# Communication networks of wild zebra finches (*Taeniopygia castanotis*)

**DOI:** 10.1101/2025.09.11.675577

**Authors:** K. Hagedoorn, N. Tschirren, E. ter Avest, C. Tyson, L. Snijders, S. C. Griffith, M. Naguib, H. Loning

**Affiliations:** Behavioural Ecology Group, Wageningen University & Research, Wageningen, The Netherlands; School of Natural Sciences, Macquarie University, Sydney, New South Wales, Australia; School of Biological, Earth and Environmental Sciences, University of New South Wales, Sydney, New South Wales, Australia

**Keywords:** Communication networks, birdsong, multi-level society, social network, zebra finch, BirdNET, fission-fusion societies, vocal communication, environmental unpredictability

## Abstract

Communication networks are widespread across species, permitting information flow and facilitating social connections across space and time. In birds, communication networks are well studied in territorial species with long-range songs connecting individuals across space, where unintended listeners extract information from others’ signalling interactions. Yet, acoustic signals also play important social roles at short range, forming communication networks that connect individuals within larger social units. Wild zebra finches (*Taeniopygia castanotis*) provide a unique model system to examine such communication networks in a non-territorial species. Zebra finches breed in loose colonies in the Australian arid zone, move around in pairs or small groups, and gather at social hotspots, thus forming dynamic, potentially multi-level, societies. Here, we quantified singing activity and connectivity using the individually distinctive male song recorded at two breeding sub-colonies and three social hotspots over one to three days. We identified 1,835 song bouts from 163 males based on spectrographic similarities and we assessed within and between individual song assignments with a deep learning model (BirdNET). We constructed communication networks based on temporal singing proximity at shared locations: social hotspots and breeding colonies. We reveal higher singing activity at social hotspots than at breeding sites, with almost no dawn song at either site. Singing peaked later, yet at different times of day between breeding sites and social hotspots. Communication networks, with distinct males singing in close temporal proximity, were apparent in both contexts, with larger networks at hotspots. These networks included some individuals that sang together repeatedly at either site, but overall networks were not strongly nested, with only very few males maintaining associations across breeding colonies and hotspots. These networks may facilitate synchronised foraging and breeding as adaptations to a harsh and unpredictable environment. Additionally, our approach offers a novel road map for widening the understanding of communication networks.

## Introduction

Animals vary in communication and sociality, ranging from simple to complex signalling interactions and from solitary living to highly complex social groupings and networks (Ward & Webster, 2016). Sociality and communication are tightly linked, yet while social networks typically refer to spatial proximity of individuals (Whitehead, 1997; Croft et al., 2008; Krause et al., 2011), communication networks have been mainly considered for long-range signalling interactions where individuals do not associate closely (McGregor, 2005). Such long-range signals enable information transfer across large distances and can thus be crucial to determine movements, mate choice, breeding decisions, territorial behaviour, responses to predators and other aspects of social synchronisation (Waser & Wiley, 1980; Naguib & Wiley, 2001; Snijders & Naguib, 2017). In many cases, these signals peak at specific times of day, such as in the dawn song of territorial songbirds which is most commonly associated with territorial defence and mate attraction (Staicer et al., 1996; Kunc et al., 2005).

Studies on such long-distance signalling have shown that information is not only gathered from a singer and its song but also by eavesdropping on the dynamics of vocal interactions among others, information that is not available when attending to singers separately, leading to the initial definition of communication networks (McGregor & Dabelsteen, 1996). Such relative differences in singing during vocal interactions can provide information for reproductive decisions (Otter et al., 1999; Schmidt et al., 2006) and affect priorities in territory defence (Peake et al., 2001; Mennill & Ratcliffe, 2004; Naguib et al., 2004; Peake et al., 2005; Fitzsimmons et al., 2008; Amy et al., 2010; Foote et al., 2010). Eavesdropping on vocal interactions among others has been at the core of communication network approaches to birdsong (Peake, 2005; Naguib et al., 2011). Yet, biologically, a communication network, defined as patterns of potential information flow, would be much larger, also including information extracted from individual songs and from direct interactions. Such wider approaches can be useful to better understand the link between communication networks and spatial networks, as well as to obtain a better understanding of information flow across large and complex social organisations in animal societies (Snijders & Naguib, 2017; Reichert & Caves, 2026).

Because information from signals affects social decision making, including the initiation, maintenance and dissolution of social bonds, signals can be expected to play a key role also in the social organisation of non-territorial animal societies. For instance, in dynamic and more complex animal organisations such as multi-level fission-fusion societies, in which not only individuals but also social units at higher levels of organisation fission and fuse (Papageorgiou et al., 2019; Papageorgiou & Farine, 2021), signals carrying individually specific information might be key facilitators allowing individuals to maintain one-to-one or pair-to-pair connections within larger social units. However, the quantitative use of individually-distinct signals by animals in dynamic, potentially multi-level, societies has received little attention (Snijders & Naguib, 2017; Reichert & Caves, 2026).

The zebra finch (*Taeniopygia castanotis*) in its natural habitat offers a highly suitable study system to address these knowledge gaps. Zebra finches are an important model organism for many lab-based studies (Griffith & Buchanan, 2010; Hauber et al., 2021), and their dynamic social organisation in the wild offers scope to broaden the traditional concept of communication networks in order to better understand the role of signals in social organisation (Loning et al., 2024). Zebra finches live throughout the Australian arid zone and move around together with their mate (Tyson et al., 2024) and are encountered mostly as a pair and not in a group (McCowan et al., 2015). These pairs gather at social hotspots, frequently visited gathering sites that likely are used to socialise, during non-breeding times (Loning et al., 2023a) and additionally associate with others in their loose breeding colonies during breeding (Brandl et al., 2021). When resources are highly clumped, birds can form temporarily large groups, such as near water (Zann, 1996; McCowan et al., 2015). Social network structure is dynamic across the day and weather conditions, with higher modularity (more sub-units) early and late in the day and at higher wind speeds (Tyson et al., 2026). These fission-fusion dynamics suggest that in the wild, zebra finches might live in multi-level societies (e.g. pairs within loose groups, loose groups within larger aggregations), in similar ways as shown for aviary conditions (Zhang et al., 2025). Since male zebra finches each sing a single unique song motif with minor variation (Hauber et al., 2010) and both sexes have individually distinct calls (Elie & Theunissen, 2018), their dynamic social organisation is most likely facilitated by these distinctive vocal signals, allowing receivers to potentially identify others in different social settings and in different locations. Zebra finch vocalisations are very short-distance signals, with an average detection range of nine meters for their song, and the loudest calls in their repertoire have an average detection range of only 14 meters (Loning et al., 2022). These individually distinct close-range vocalisations therefore likely function to establish and maintain social connections through information exchange at short range. This can include information on individual condition and foraging success (Ritschard & Brumm, 2012) and breeding state (Loning et al., 2023b) at gathering points, such as at the breeding area (Brandl et al., 2019c), at foraging sites (Brandl et al., 2019a; Brandl et al., 2021) or at the social hotspots (Loning et al., 2023a).

We took advantage of the wild zebra finch social system, in which birds vocalise and encounter conspecifics at different locations, to integrate communication networks and spatial networks (Snijders & Naguib, 2017; Loning et al., 2024) and so examine the network structure of a highly dynamic wild bird society. We quantified male singing activity throughout the day in a wild population of zebra finches in the Australian Outback, recording singing both at breeding sites and social hotspots (Loning et al., 2023a) using automated audio recorders. We used the individually distinctive male song (Sossinka & Böhner, 1980; Hauber et al., 2010), as a proxy for presence at nests and hotspots and determined if different males repeatedly sang at the same time at these locations. From these songs, we created communication networks and expected larger networks at the hotspots, with more individuals heard singing and associating there than in the breeding colonies. We also expected a hierarchical social structure with individuals at the hotspots interacting with conspecifics they associated with in the breeding colony, as well as others. Although only males sing, leading here to male-focussed networks, pairs stay almost constantly together (Tyson et al., 2024) and typically two individuals arrive and leave the hotspots together (Loning et al., 2023a). Pairs also typically arrive together at their nestbox when breeding (Mariette & Griffith, 2012b), suggesting that male song networks in most cases most likely reflect a network among pairs.

## Methods

The study was conducted from September to October 2024 in the Gap Hills area of the Fowlers Gap Arid Zone Research Station, New South Wales, Australia. This area consists of open shrubland with low bushes such as bluebush (*Maireana* sp.) and saltbush (*Rhagodia* sp.) as well as higher vegetation along natural creeks, mostly shrub-like trees such as prickly wattle (*Acacia victoriae*) and dead finish (*Acacia tetragonophylla*). There is an artificial water reservoir in the area (Fig. 1), and zebra finches at this site use nestboxes to breed, which are placed on poles near bushes or trees for shelter (Griffith et al., 2008). Nestboxes were placed in the area in six sub-colonies of about 30 boxes each.

**Figure 1.**
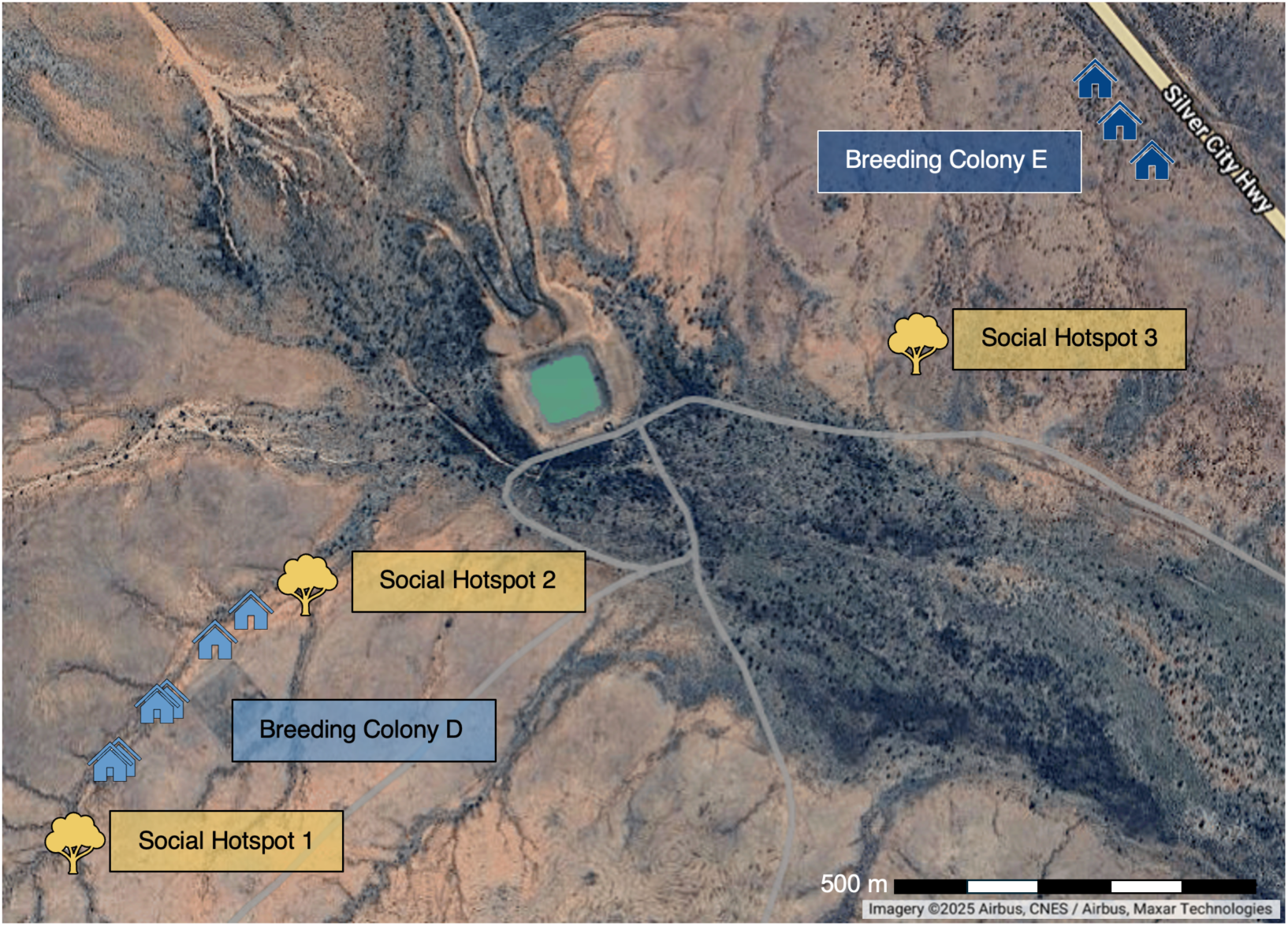
Map showing the field site at Gap Hills with positions of analysed nestboxes and social hotspots. Tree icons denote social hotspots, house icons denote nestboxes selected for audio analysis, with six nestboxes located in breeding colony D and three nestboxes located in breeding colony E. The central dam, creek lines and highway are visible. Satellite image retrieved from Google Maps.

### Audio data collection

We recorded vocal activity at nestboxes and nearby social hotspots using time-programmable audio recorders (AudioMoth; Hill et al., 2019). AudioMoths were set to record 16-bit wav files from 90 minutes before sunrise to 90 minutes after sunset (settings: 32 kHz sample rate; medium gain; energy saver mode on). We monitored breeding activity by checking all nestboxes in the area every three days and attached AudioMoths to nestbox poles of active nests. During the study period, we observed a total of 103 breeding attempts (egg-laying stage or beyond) in 78 boxes. Most breeding attempts occurred in breeding colonies ‘D’ and ‘E’ (Fig. 1) with 38 attempts in each area in 28 and 27 boxes, respectively. We identified social hotspot locations by walking repeated transects, registering recurring observations of zebra finches and checking for the presence of faecal droppings. We combined this information with hotspot locations obtained in the same way in previous years, as social hotspots are generally temporally stable (Loning et al., 2023a). We selected three out of 13 potential social hotspots and observed these three potential hotspots three times for one hour between 0800 and 1230 hours on different days to confirm that they were actively used by zebra finches. We then placed AudioMoths at these three identified social hotspots (all three were also in use in the previous year). Two of these social hotspots were located at the edges of a breeding sub-colony (colony D), and one was located between another sub-colony (colony E) and the central water reservoir (Fig. 1).

### Audio analysis

For each of nine nestboxes (Fig. 1), we selected one day of audio recordings for analysis, as one day has been shown to be sufficient to record the locally breeding birds (Loning et al., 2023b). We selected the day with the most favourable weather conditions (i.e. no rain, little wind) during the egg-laying phase as males sing most during this phase of breeding (Loning et al., 2023b). The selected nests all produced fledglings but were active at different times during the study period. For the three hotspots, we selected audio recordings from seven different days, two days for hotspots 1 and 2 and three days for hotspot 3 (Fig. 1). Where possible, we analysed recordings from hotspots on the same day as a nearby nestbox (at most five days difference), to minimise environmental variation such as wind affecting movement and sociality (Tyson et al., 2026). In total, we analysed 248 hours of audio recordings, consisting of 140 hours of recordings at nine nestboxes (one day per nestbox) and 108 hours at three social hotspots (two to three days per hotspot).

Within these audio recordings, we first assigned zebra finch songs to individual males by listening to the song recordings and visually inspecting their sound spectrograms (window type: Hamming, window size: 1024, window overlap (hard-coded): 50%, gain: 20 dB, dynamic range: 80 dB) in Audacity (version 3.7.1; Audacity Team, 2024). Zebra finch song has a distinct structure, with each male repeatedly singing an individually distinct motif (series of syllables), which is readily recognisable by conspecifics (Hauber et al., 2010) as well as by human biologists using spectrograms (Fig. 2) although some intraindividual variation exists (Sossinka & Böhner, 1980; Sturdy et al., 1999). A gap of more than five seconds between song motifs separated song bouts (Zann, 1996). We assigned a recording quality score to each song bout on a scale of one (song hardly audible and only the loudest distance call elements are visible on the spectrogram) to ten (clear, unmasked spectrogram). Only songs of high quality, with a score of six (song is audible but distant, there is some masking, spectrogram mainly clear) or higher, were assigned to an individual bird, while songs of lower quality were not included here. We established a database containing clear spectrograms of individual birds and compared newly encountered songs to this database. If the shape of the frequency spectrum of song elements, their sequence within motifs, as well as the motif organisation of a new song, clearly matched that of an already encountered individual, the observer assigned it to the same individual. Ambiguous cases were always checked by a second observer. If there was no consensus on the classification, that song was not assigned to a specific individual. Out of the total 2,552 song bouts present in our recordings (248 hours), we assigned 1,835 song bouts (71% of song bouts) to 163 individuals (mean ± SD: 11.3 ± 15.9 song bouts per individual, range 1-113).

**Figure 2.**
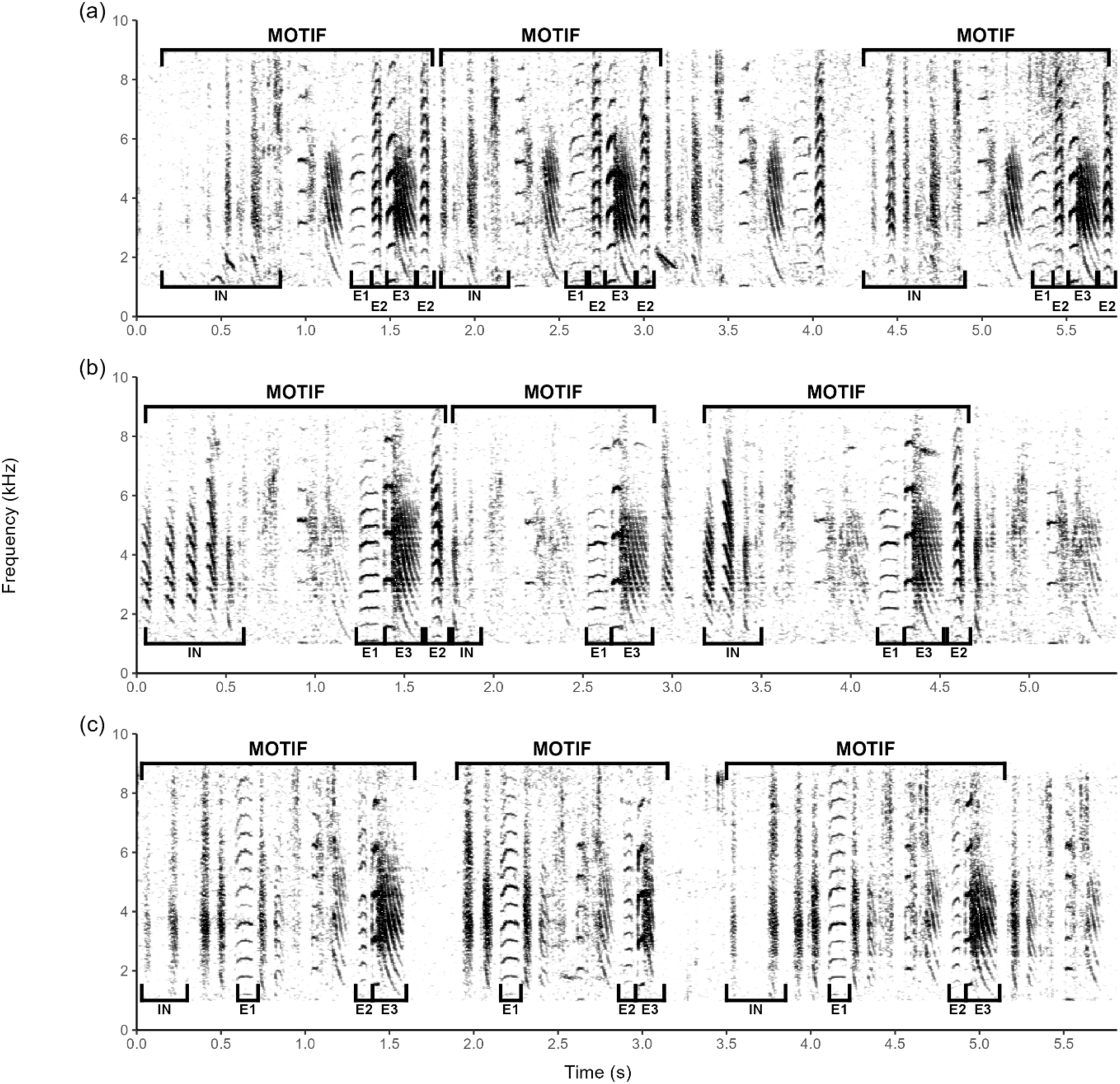
Example spectrograms of song bouts from three different individuals: (a), (b) and (c). The y axis shows the frequency in kHz and the x-axis Ume in seconds. Darker shades correspond to a higher sound energy. Different elements of the moUfs are annotated. IN denotes introductory notes, E1, E2 and E3 denote different song element types. The three individuals (a-c) differ in element sequence, and elements of the same type differ in shape of frequency spectrum as well as fundamental (and therefore, harmonic) frequency between individuals. Intraindividual differences such as variaUon in moUf duraUon and content are visible, e.g. individual 2 does not have element E2 in the second moUf. Spectrograms were made using the ‘seewave’ package in R (window size: 1024, window type: Hamming, window overlap: 80%, frequency limit: 1-9 kHz, amplitude 204 normalised, only bins in the fourth quarUle are plojed).

Using all 2,552 song bouts, we visually compared song activity over the day at both location types (nestboxes: *N* = 9 days; social hotspots; *N* = 7 days) by summing the number of seconds containing song within each half-hour block, (expressed as percentage of all daily song activity). Additionally, we compared the total daily song activity (in minutes of song per day) between nestboxes and social hotspots using all song bouts. Using the 1,835 song bouts that were assigned to individuals, we compared the daily number of unique individuals singing at both location types.

### BirdNET validation of audio analysis

Although the distinctive individual signature of zebra finch song is well established, identifiable also in visual inspection of spectrograms, (Immelmann, 1968; Böhner, 1983; Zann, 1996; Hauber et al., 2010; Woodgate et al., 2012), we additionally tested if within and between individual differences in songs from our individual assignments were also reflected in a machine learning approach. Therefore, we used BirdNET (model version 2.4; (Kahl et al., 2021)) as an impartial observer to validate whether or not the songs we had manually assigned to an individual based on visual inspections of sound spectrograms were indeed more similar to each other than to songs that we assigned to other individuals, while accounting for unavoidable variation in recording conditions. BirdNET is a deep-learning model trained to recognise the vocalisations of over 6,000 bird species to the species level (including zebra finches), but it has not been trained to distinguish between zebra finch individuals. BirdNET was used as a pre-trained model without additional training or fine-tuning on our dataset, as this would have required using our individual labels, which we intended to validate independently. BirdNET processes each sound fragment and extracts a set of 1,024 numerical features. These so-called embeddings are a compact representation that captures the patterns relevant for distinguishing bird species (McGinn et al., 2023). While originally designed to support species classification, the embeddings can also be treated as a 1,024-dimensional feature vector that characterises the acoustic properties of individual bird sound recordings.

We extracted BirdNET embeddings representing zebra finch songs for subsequent intra- and inter-individual comparison. Since BirdNET internally uses three-second segments for classification, we trimmed identified song bouts (*N* = 1,835) to the first three seconds and also kept the last 3 seconds for bouts over 6 seconds long. Song bouts shorter than three seconds were left unaltered (BirdNET pads segments shorter than three seconds with trailing zeroes). This resulted in 2,617 song segments (belonging to 163 individuals) for which we extracted the corresponding embeddings using BirdNET-Analyzer (GUI version 2.1.1; settings: overlap 0 seconds, batch size 1, no speed modification, bandpass frequency 0-15 kHz). We then calculated cosine distances (i.e. 1 - cosine similarity, which measures the angle between vectors irrespective of their magnitude) between the embeddings for all combinations of segments using SciPy’s *spatial.distance.pdist* (with metric=“cosine”) function in Python (version 3.13.5). This resulted in a combined 3.4 million intra- and interindividual comparisons. We further divided comparisons into three categories based on where the two compared segments originated: comparisons made between segments from (1) the same one-hour audio recording, (2) the same date, and (3) different dates. These categories reflect a potential increase in recording condition differences and a decrease in certainty of songs actually having been sung by the same individual.

### Video recordings at social hotspots and analysis

To quantify zebra finch visit durations for constructing communication networks, we filmed the three social hotspot trees using tripod-mounted action cameras (GoPro Hero5 Black, resolution of 1280×720, 25 FPS, narrow field of view) for a total of six days (one to three days per social hotspot) from a height of about 150 cm at a distance that allowed the whole tree to be in frame. To estimate an average time zebra finches spend at a social hotspot, we analysed 40.2 hours of video footage of the three social hotspots (mean ± SD: 6.71 ± 2.64 hours per day, range 3 – 9.7 hours). Each departure from and arrival at the hotspot was scored using BORIS (Behavioural Observation Research Interactive Software; version 9.2.1). Then, per day, we matched each arrival to the earliest unmatched departure to calculate the duration birds spent on the tree (Fig. 3). To compute these visit durations, we excluded visits lacking matching pairs attributable to missed arrivals or departures on video and birds staying for extended periods. To limit the influence of extreme outliers and to focus on typical visit durations, bird visits lasting longer than one hour (*N* = 45 visits) were excluded from the main analysis. This cut-off was based on preliminary data exploration, which showed that durations exceeding one hour were rare and probably represented atypical or faulty events. Applying this filter improved the robustness of summary statistics while retaining the majority of observations. This resulted in 686 bird visits with a mean hotspot visit duration of 1,263 seconds ± 722 s (± SD; median: 1,364 seconds; IQR 634 – 1,750 seconds).

**Figure 3.**
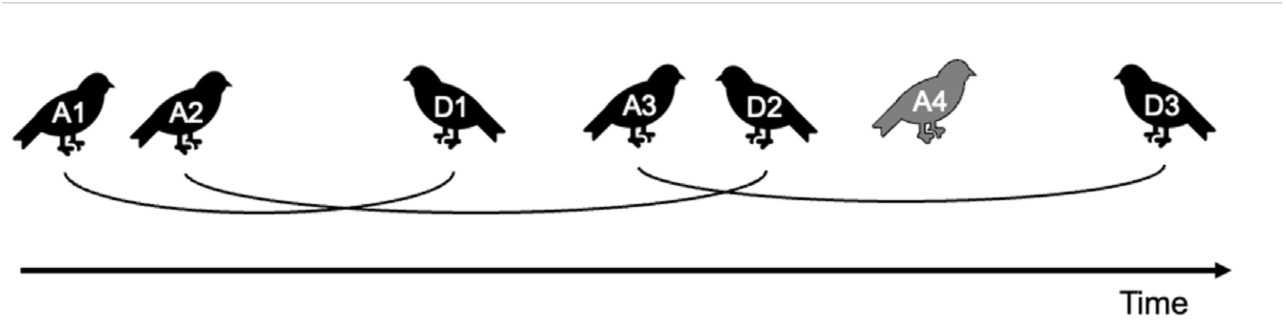
Schematic overview of matching up each arrival (A) and departure (D) scored in the video analysis to measure visit duration of birds to a social hotspot. From the start of the day, each arrival was matched to the next possible departure, i.e. the first arrival (A1) was matched to the first departure (D1), the second arrival (A2) was matched to the second departure (D2), with all subsequent arrivals paired to departures in the same order. Unmatched arrivals or departures (such as A4) were left unmeasured.

### Communication network characteristics

We determined network edges (associations) on the assumption that individuals singing together were associating. Wild zebra finches sing softly (Loning et al., 2022). Therefore, recorded songs that were clear enough to be individually distinguished by us were likely produced by birds perched in the bush next to each nestbox or the social hotspot itself. Given the average detection range for zebra finch song of nine meters (Loning et al., 2022) and the sparse distribution of vegetation in our study area, birds within the same bush (nestbox or hotspot) would be within this hearing range, whereas birds on the next bush would likely not.

In order to determine individual associations, we first established individual visits based on identified song bouts. We combined all consecutive song bouts of an individual that fell within 1,263 s of each other (the mean visit duration at social hotspots, see above) into a single visit. This resulted in 418 visits with an average duration of 324 ± 581 s (± SD; median: 52 s; IQR: 9.5 – 289 s; range 1.4 – 3,475 s). We then considered individuals that had overlapping visits to be associating and constructed three networks (Fig. 4), which differed in assumed visit duration. In the most conservative network (referred to as ‘Conservative network’ from here on), no extra assumptions regarding visit duration were made, and we considered birds to be present from the start of their first song bout until the end of their last song bout. However, it is unlikely that birds attending a nestbox or social hotspot are present exclusively for the duration of their singing activity (pers. obs.). Therefore, for the other two networks, we assumed that birds would visit at least for the duration of 634 s (the first quartile visit duration as quantified from video recordings, see above; the ‘Q1 network’) or 1,263 s (the mean visit duration; the ‘Mean network’), measured from the start of the first song bout. For each network per location type (nestbox, social hotspot, both), we calculated the number of associating individuals that were singing there (nodes), the number of associations (edges), and the number of dyads. We also calculated the number of dyads with one, two, or more associations as well as the network density.

**Figure 4.**
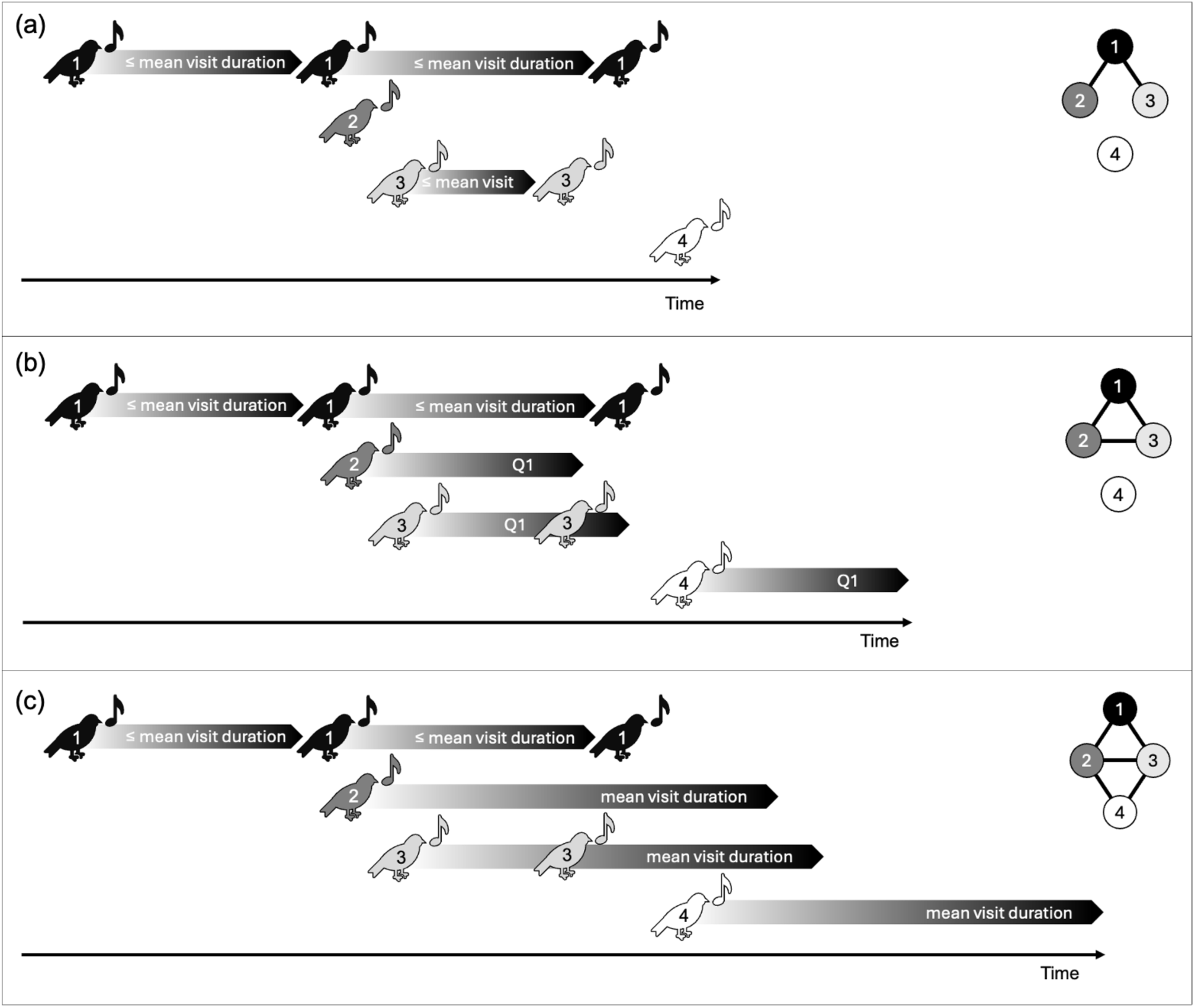
Schematic overview of the three constructed networks based on identified song bout visits. Each song bout is illustrated with a pictogram of a singing bird, and individual birds are numbered accordingly. For all networks, visits started at the first song bout and included all subsequent song bouts occurring within the mean visit duration of each other. Song bouts with gaps shorter than the mean visit duration were combined, so actual visits could exceed the mean duration calculated from the video analysis. In the most conservative approach (a) Conservative network, a visit ended at the end of the last song bout, meaning that birds then only sang once were considered to only visit for the duration of that song bout. As birds likely do not only visit for the duration of singing, we also constructed slightly less conservative networks: (b) Q1 network, visits that in the Conservative network were shorter than the first quartile visit duration (Q1) were extended to Q1, whereas visits already exceeding Q1 (such as bird 1) remained unchanged. (c) Mean network, all visits shorter than the mean visit duration were extended to the mean visit duration. Birds that had a total visit duration already exceeding the mean visit duration (again bird 1) remained unchanged. Diagrams on the right show the resulting network with circles representing birds (nodes) and connecting lines as song-based associations (edges).

We further assessed the potential for associations beyond a dyadic nature, *i.e.* triads and larger groups. To do so, we performed a sliding window analysis. We constructed hypothetical visits with a duration of 1,263 s (the mean visit duration), with start time differences of one minute to quantify the distribution of encountered identified singing individuals in those time windows. Before calculating summary statistics, we filtered the time windows to contain at least one singing individual to avoid parts of the day in which zebra finches did not sing and thus could not associate anyway in this dataset.

For individuals that were heard both at a nestbox and a social hotspot (*N* = 24), we calculated the number of associations and the number of unique associations (i.e. the number of unique individuals associated with) they had at nestboxes and social hotspots. If the bird was heard at multiple nestboxes or multiple social hotspots, we used the total number of associations or the number of unique associations and calculated how long each individual sang (summed duration of all the song bouts) at each type of location.

### Statistical analysis

All statistical analyses were performed using R (version 4.5.1; R Core Team, 2025). Linear mixed-effect models (LMMs) and generalised linear mixed-effect models (GLMMs) were constructed using the *lmer* and *glmer* functions from the ‘lme4’ package, respectively (Bates et al., 2015). Model assumptions were checked using the *simulateResiduals* and *testDispersion* functions (for Poisson GLMMs) from the ‘DHARMa’ package (Hartig, 2024). We tested for statistical significance of variables with two categories using estimated marginal means (EMM) with the *emmeans* function from the ‘emmeans’ package (Lenth, 2025). We also used this function to obtain the reported mean ± SE and contrasts for all mixed model data. To test interaction effects and variables with more than two categories, we compared an interaction model to a model without the interaction term with a likelihood-ratio test, using the function *anova* from the ‘stats’ package.

To test for differences in singing at the nestboxes and social hotspots, we compared the total duration of song and the number of individuals heard per day between social hotspots and nestboxes using a LMM and a GLMM with a Poisson distribution, respectively. For both models, we accounted for the different number of observation days for the different locations (*N* = 1 for all nestboxes, *N* = 2 for hotspot 1 and 2, *N* = 3 for hotspot 3) by adding the location as random intercept.

For each comparison between BirdNET embeddings, we calculated the cosine distance. We then used a LMM to test the effect of comparison type (intra- or interindividual) as well as song origin (from the same one-hour audio recording, the same date, or different dates) and their interaction on cosine distance. We included the assigned individuals of both compared songs (the individual for song 1 and for song 2 was the same in the case of an intraindividual comparison) as crossed random intercepts.

To quantify the communication/social networks, we obtained network metrics and graphs using the *igraph* package (Csárdi & Nepusz, 2006). For the nestboxes and social hotspots, we calculated and compared the number of nodes (order) and edges (size) of the repeated association network (defined as dyads with two or more interactions between them) per day using Poisson GLMMs with location as random effect. We then compared the total number of associations per individual and the number of unique associations (dyads) between breeding sites and social hotspots using Poisson GLMMs with individual ID as a random intercept. For these models, we included song duration in minutes as a covariate. We used the Mean network for this individual-level comparison to obtain a sufficiently large sample of individuals that associated at both location types.

### Ethical note

We collected all audio and video data using non-invasive methods. Audio and video equipment was placed while minimising disturbance. Approval was granted by the Macquarie University Animal Ethics Committee, and the study was conducted under a New South Wales DPIE Scientific Licence SL100378, Ethics ARA 2018/02.

## Results

### Singing activity at breeding sites and social hotspots

In 246 hours of recordings, comprising nine nestboxes (breeding sites) at two breeding sub-colonies and three hotspots, a total of 163 males were identified individually by their song. Most individuals were heard at only one location (*N* = 126), with fewer individuals heard at more than one location (two locations: *N* = 23, three locations: *N* = 9, four locations: *N* = 4, five locations: *N* = 1). In total, 24 individuals (14.7%) were heard at both a social hotspot and the breeding sites. Zebra finch males hardly sang at dawn at either location (Fig. 5a). Almost all birds started singing after sunrise later in the day around the same time of the day at both locations, breeding sites and social hotspots (Fig. 5a). Yet, at the breeding sites, most song occurred earlier in the day between sunrise and noon. At the social hotspots, most song occurred later in the morning and early afternoon. Furthermore, song continued at the social hotspots for longer than at the breeding sites. The average duration of song in minutes per day at social hotspots (mean ± SE: 23.8 ± 4.3 min, *N* = 7) was significantly higher than at breeding sites (6.8 ± 3.3 min, *N* = 9; EMM: *t*(5.9) = 3.2, *P* = 0.02; Fig. 5b). Furthermore, significantly more individuals were heard during the day at social hotspots (20.8 ± 4.9 individuals, *N* = 7) than at the breeding sites (8.9 ± 1.5 individuals, *N* = 9; EMM: *z* = 2.9, *P* = 0.004; Fig. 5c).

**Figure 5.**
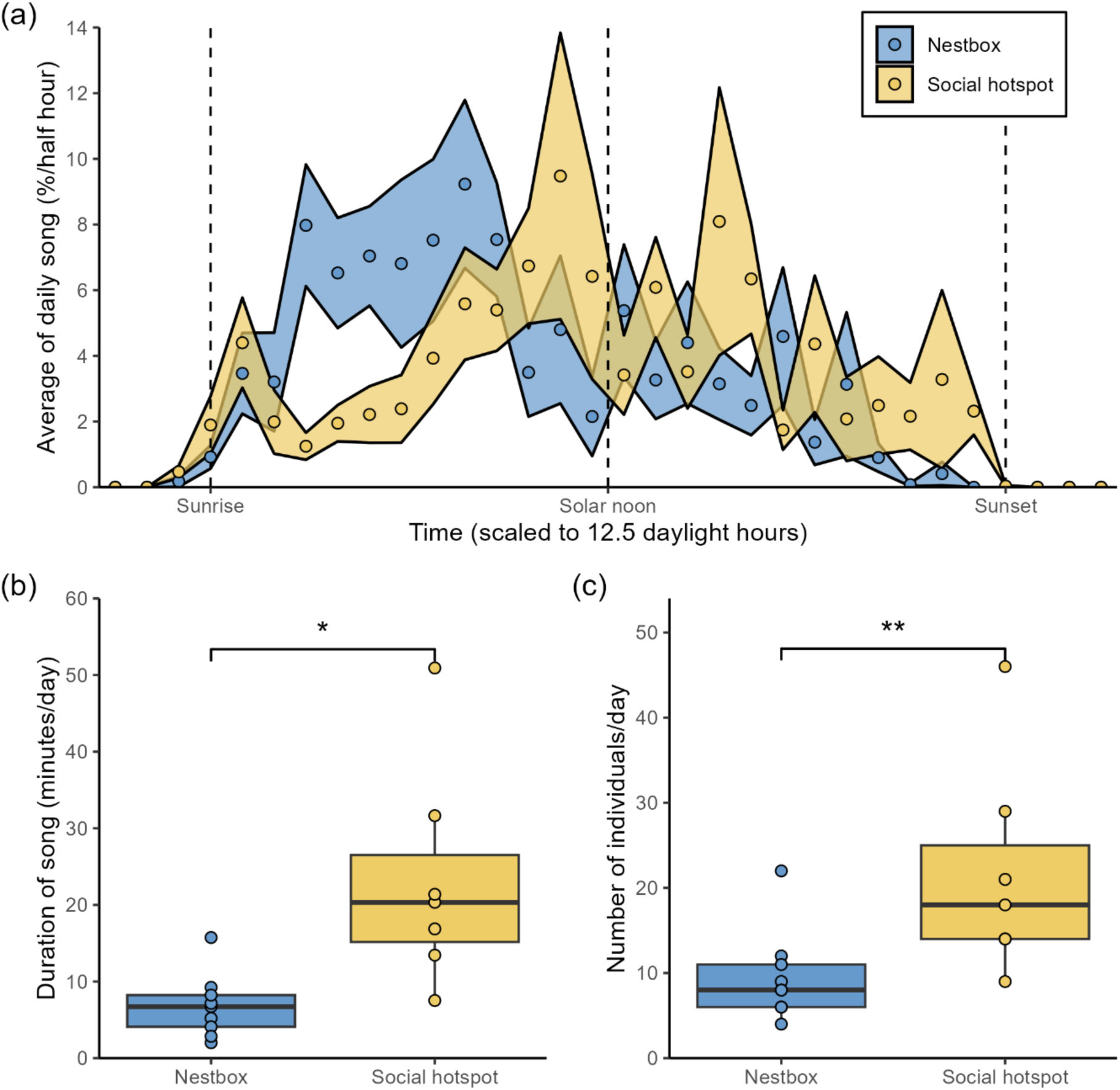
(a) The average distribution (points) ± SE (polygons) of song per half hour over the day for breeding sites (nestboxes, n = 9 days) and social hotspots (n = 7 days). The time is scaled to an average day (sunrise to sunset) of 12.5 hours. Sunrise, solar noon, and sunset are denoted by the dashed vertical lines. The y-axis shows the proportion of total daily song that occurred per half hour. The average duration of song in minutes per day (b) and the number of individuals heard per day (c) was higher at social hotspots than at breeding sites. * denotes *P* < 0.05, ** denotes *P* < 0.01.

### BirdNET validation of song classification

Songs that we assigned by visual inspection of sound spectrograms as produced by the same individual were more similar using BirdNET analyses, as cosine distance for intra-individual song comparisons (0.304 ± 0.004) was significantly lower than for inter-individual comparisons (0.362 ± 0.004; EMM: *z* = 156.3, *P* < 0.001; Fig. 6). There was also a significant interaction between song comparison type (same individual and different individual) and sound origin (same bout, same day, different day) (likelihood-ratio test: χ^2^(2) = 578.5, *P* < 0.0001). The difference between intra-individual and inter-individual song comparisons was largest for compared songs from the same one-hour recording (0.070 ± 0.001), smaller for the same date (0.053 ± 0.001), and smallest for different dates (0.050 ± 0.001; all *P* < 0.0001; Fig. 6). However, the BirdNET embeddings probably did not exclusively reflect zebra finch song, but also reflected recording conditions, as sound origin differences also affected the cosine distance (Fig. 6). Overall, songs originating from the same one-hour recording had lower cosine distances (0.305 ± 0.004) than those recorded at the same date (0.340 ± 0.004), and these in turn were lower than those comparisons with songs recorded on different dates (0.354 ± 0.004; likelihood-ratio test: χ^2^(2) = 19,346, *P* < 0.0001). These differences likely cannot be attributed exclusively to potential errors in individual assignment as not only the intra-individual, but also the inter-individual comparisons originating from the same one-hour recording had lower cosine distances (intra: 0.270 ± 0.004; inter: 0.340 ± 0.004) than those made between recordings of the same date (intra: 0.313 ± 0.004; inter: 0.366 ± 0.004) and those of different dates (intra: 0.329 ± 0.004; inter: 0.380 ± 0.004; EMM: all pairwise *z* > 16.7 and *P* < 0.0001; Fig. 6).

**Figure 6.**
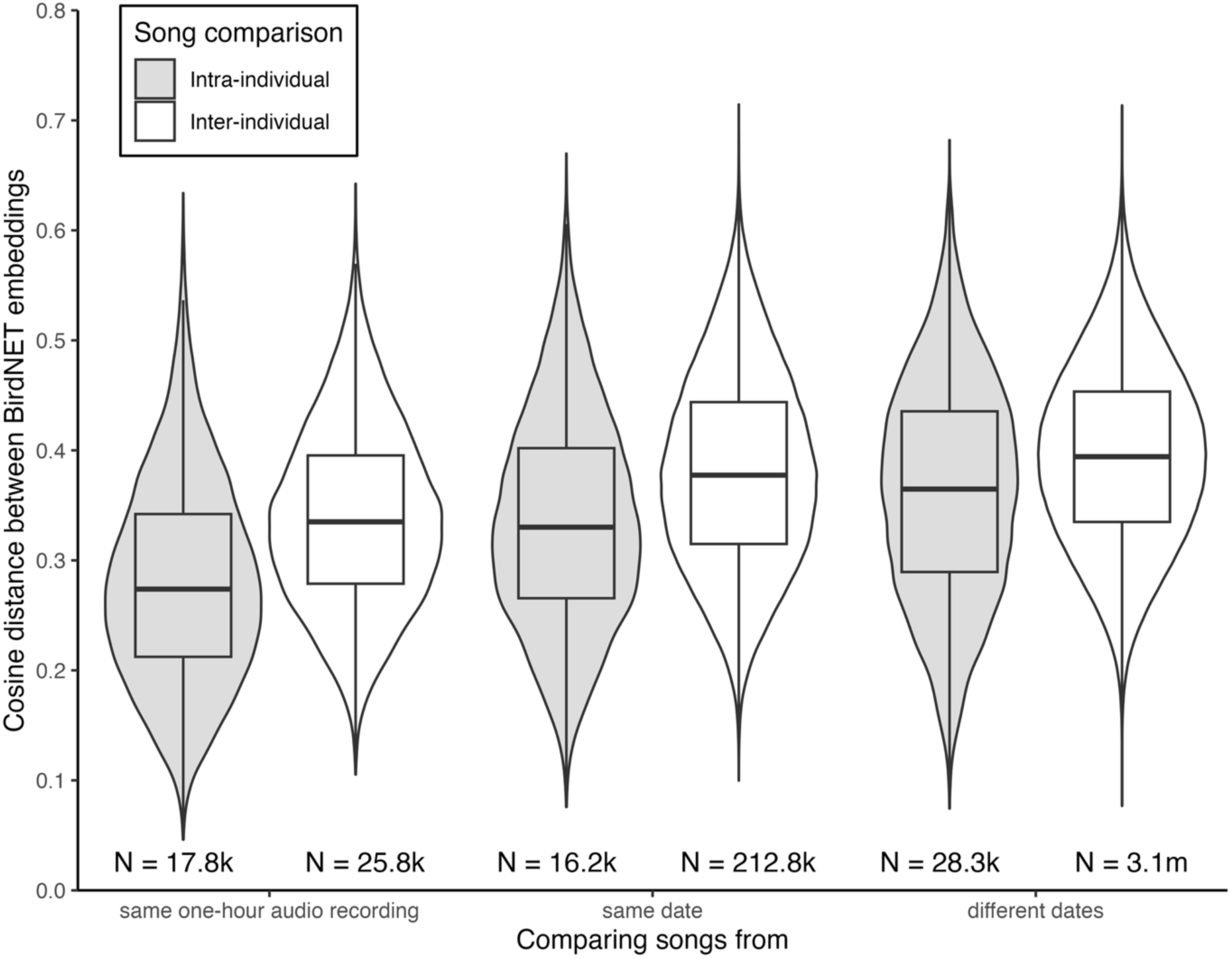
Comparisons of extracted BirdNET embeddings as validation of individual identification. We calculated cosine distances between embeddings of 2,617 songs, resulting in 3.4 million comparisons (Sample size per comparison is given on the x-axis, k = thousand, m = million). The cosine distance or dissimilarity ranges from zero (vectors have the same direction) to two (vectors have the opposite direction). Across song origins, cosine distance was lower for intraindividual song comparisons (i.e. between songs of the same individual as assigned based on spectrographic similarities) compared to interindividual comparisons. This difference was largest when recording conditions were similar (i.e. with both songs originating from the same one-hour audio recording), whereas it was smallest when conditions were potentially most different (songs recorded on different days). Each song comparison type-origin combination differed from the other five (EMM: all pairwise *z* > 16.7 and *P* < 0.0001).

### Communication network characteristics

In all constructed networks, zebra finches showed non-uniform network structures at both breeding sites and social hotspots. The Conservative network, Q1 network, and Mean network differed quantitatively, increasing in number of associating individuals, associations and dyads (unique associations) with increased assumed visit durations, but qualitatively the network structure of the networks remained similar (Table 1, Fig. 7). A large portion of the 163 identified individuals had an association (Conservative network: 76 individuals, 47%; Fig 7a; Q1 network: 127 individuals, 73%; Mean network: 149 individuals, 91%). However, most of the observed dyads consisted of only one association (Conservative network: 95%; Q1 network: 94%; Mean network: 89%), while only a small fraction of dyads associated three or more times (1-3%; Table 1; Fig. 7a). In the Mean network, three out of 36 dyads with two or more associations between them associated at both location types (Fig. 7b), indicating that most individuals, within the analysed time frame, had repeated social ties either at the breeding site or at social hotspots, but not both. For all three network types (Conservative network, Q1 network, Mean network), the network on a given day had more nodes and more edges at social hotspots than at breeding sites (Table 2).

**Figure 7.**
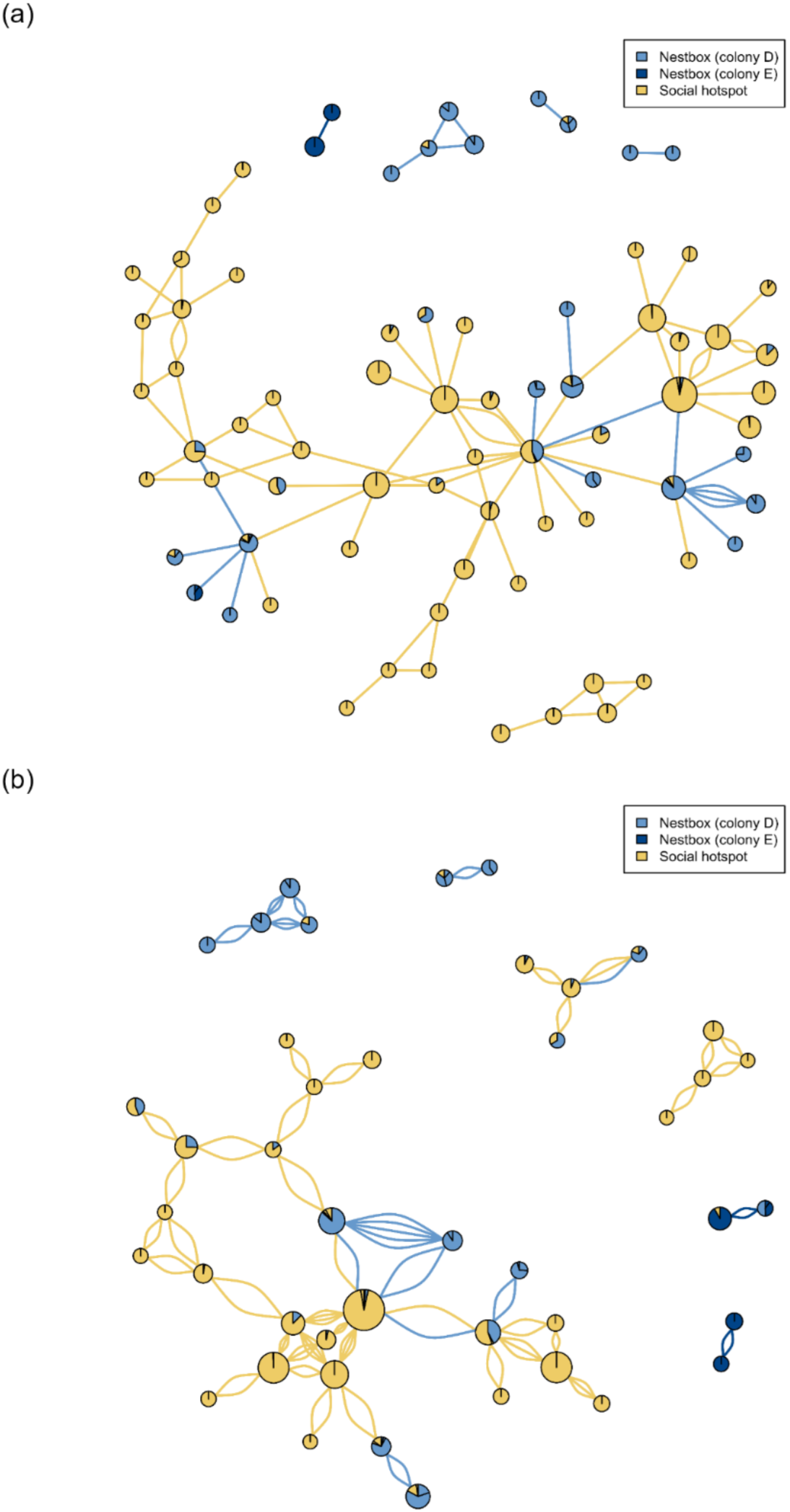
The social network for (a) all associations of the Conservative network and (b) all repeated associations of the Mean network, i.e. dyads that were identified together at least two times. The pie shapes represent associating individuals (nodes), with coloured slices representing the relative amount of time they spent singing in different locations. The size of the node scales by the total amount of time that node was heard singing. The lines represent the song-based associations (edges). The colours of the edges correspond to the location where that association took place.

**Table 1.** Network metrics of the three constructed networks based on identified song bout visits. Individuals associated when their visits overlapped. In all three networks, individual visits were calculated as starting from the first observed song bout of a given individual and included all consecutive song bouts that fell within the mean visit duration of each other (1,263 s, as observed from 40 hours of video of social hotspots). The networks differed in the assumed visit duration, with visits of the most conservative network ending at the end of their last song bout, whereas the other two networks assumed individuals would stay for at least the first quartile (634 s) or mean visit duration (1263 s), respectively. Although the networks differ quantitatively, they are qualitatively similar with most dyads consisting of only one association, and more associations and dyads at hotspots than at breeding sites. There were no or only few dyads that associated at different location types (this was observed only in the Mean network). Network densities (realised edges/total possible edges) were similar in scale.

| Network type | Associating individuals (nodes) singing at |  |  | Associations (edges) at |  | Dyads that were at |  |  | Dyads with n associations |  |  | Density |
| --- | --- | --- | --- | --- | --- | --- | --- | --- | --- | --- | --- | --- |
|  | Nest | Hotspot | Both | Nest | Hotspot | Nest | Hotspot | Both | 1 | 2 | 3+ |  |
| Conservative | 19 | 51 | 6 | 22 | 77 | 19 | 73 | 0 | 87 | 4 | 1 | 0.03474 |
| Q1 (visits 634+ s) | 26 | 81 | 20 | 60 | 207 | 53 | 191 | 0 | 230 | 9 | 5 | 0.03337 |
| Mean (visits 1263+ s) | 34 | 94 | 21 | 108 | 379 | 89 | 332 | 3 | 378 | 34 | 12 | 0.04417 |

**Table 2.** Estimated marginal means and pairwise contrasts of Poisson GLMMs comparing order (number of nodes) and size (number of edges) of daily networks between nests (n = 9) and social hotspots (n = 7). Recording location was added as a random effect.

| Daily network type | Variable | Nest | SE | Hotspots | SE | contrast ratio | SE | z-ratio | P value |
| --- | --- | --- | --- | --- | --- | --- | --- | --- | --- |
| Conservative | Order | 2.41 | 1.08 | 12.09 | 7.48 | 0.199 | 0.15 | 2.12 | 0.034 |
|  | Size | 1.63 | 0.7222 | 8.22 | 4.68 | 0.198 | 0.14 | 2.26 | 0.024 |
| Q1 | Order | 8.48 | 1.96 | 32.12 | 11.4 | 0.264 | 0.11 | 3.16 | 0.002 |
|  | Size | 5.62 | 1.49 | 23.83 | 9.39 | 0.236 | 0.11 | 3.05 | 0.002 |
| Mean | Order | 12.7 | 3.17 | 52.6 | 21 | 0.242 | 0.11 | 3.03 | 0.003 |
|  | Size | 9.79 | 2.57 | 43.44 | 18 | 0.225 | 0.11 | 3.05 | 0.002 |

The hypothetical visit windows with a mean duration (1,263 s) that included identified individuals, had on average 1.4 ± 0.7 (± SD; IQR: 1-2; range: 1-4 individuals) individuals singing at a breeding site, whereas this was 2.0 ± 1.2 (IQR: 1-2; range: 1-7 individuals) individuals at social hotspots. Of those hypothetical visits, 8% and 28% included triads or larger groups at breeding sites and social hotspots, respectively.

### Network at the individual level

In total, 24 individuals were recorded singing at both a social hotspot and a breeding site. Birds that sang longer had more associations at both location types (but this relation was not significantly different between location types; likelihood-ratio test: χ^2^(1) = 2.9, *P* = 0.09; Fig. 8a). Furthermore, these individuals had more associations at social hotspots than at breeding sites (social hotspot = 7.3 ± 1.0, breeding site = 3.6 ± 0.6; EMM*: z = 5.8, P* < 0.0001; Fig. 8b). Individuals also had more unique associations (i.e. associated with more individuals) at social hotspots than at breeding sites (social hotspot = 6.2 ± 0.9, breeding site = 3.2 ± 0.5; EMM: *z = 5.2, P* < 0.0001; Fig. 8c).

**Figure 8.**
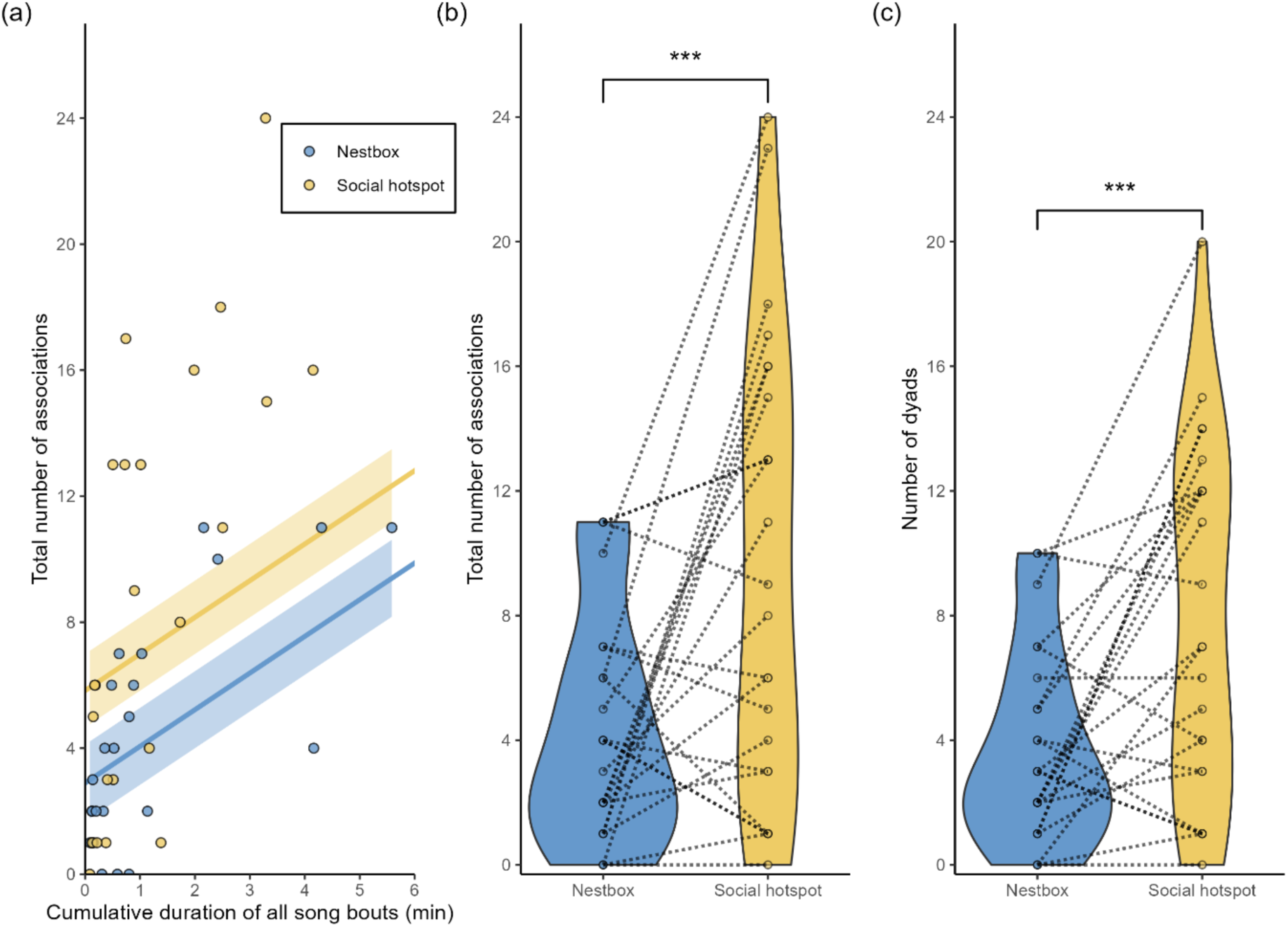
Total number of associations and number of unique associations at the individual level. Only individuals that were heard at both a breeding site and a social hotspot are shown. (a) Birds that sang more had more associations, and the number of associations was higher at social hotspots than at nestboxes. Regression lines (estimates) and shading (SE) based on the same mixed model as presented in (b) with individual as random intercept. (b) Individuals had more associations at the social hotspot than at the breeding site. (c) Individuals met with more unique conspecifics (were in more dyads) at the social hotspot than at the breeding site. *** denotes *P* = 0.001.

## Discussion

Here we reveal distinct locations at which wild zebra finches sing in close spatial and temporal proximity, namely their breeding colony and at social hotspots. Song at these locations was almost never produced at dawn, in contrast to many territorial species. Instead, it peaked after sunrise, at the breeding colony later in the morning and at the social hotspots just before noon and in the early afternoon. At both locations and in all three different analytical network approaches, multiple individually identified birds were singing in close temporal proximity in a communication network. The networks at the breeding colony and social hotspots were distinct, with networks at the social hotspots being larger than those in the breeding colony. Most individuals were heard at one location type only, and most dyads of males were heard together once. Yet, we also identified males that sang together at different times. Most dyads that associated in the breeding colony were not heard together at a social hotspot and vice versa, however a few dyads (three in the Mean network) were identified at multiple location types. These findings suggest that within this relatively short time frame studied, breeding sites and social hotspots served as distinct sites for social interactions, with a few individuals and dyads forming a bridge between them. These findings also suggest that the males of neighbouring breeding pairs do not move together as a unit to these hotspots, which can be hundreds of meters away from their breeding site. Instead, the decision to visit social hotspots is likely more a decision taken by an individual or a pair.

The higher singing activity at the social hotspots was predicted, as social hotspots are defined as gathering places (Loning et al., 2023a) and zebra finch males sing all year round at various locations (Loning et al., 2023b). Singing in different social contexts is interesting as it might be triggered mechanistically through different internal and external processes (Waas et al., 2005; Wang et al., 2012), while also having different functions and implications. Singing at the breeding colony may primarily reinforce pair bonds, play a role in nesting behaviour (Dunn & Zann, 1996), and indicate local breeding activity. Here song provides reproductive cues to neighbours (Loning et al., 2023b), which are known to coordinate their breeding with one another (Brandl et al., 2019c). As shown earlier (Loning et al., 2023b), we regularly observed many more singing individuals than just the expected resident male at a given nest site, and here we show that individuals regularly sang in close temporal proximity there. Singing at social hotspots, or in similar contexts in other species where individuals substantially vocalise in groups forming similar types of communication networks (Cure et al., 2009; Major & Jones 2011; Calcari et al., 2021; Devogel et al., 2024) might have wider social implications. Signalling in such contexts provides information to larger groups, potentially transferring information about resources, predation risk, condition and reproductive states and allowing individuals to create social bonds and perhaps coordinate breeding at a broader spatial scale.

The (almost) absence of dawn song at either location highlights an important distinction to the daily singing activity of well-studied territorial songbirds in the Northern hemisphere, where song peaks at dawn and is strongly associated with territory defence and mate choice (Catchpole & Slater, 2008). These functions appear only marginally relevant in species like zebra finches, as zebra finches are not territorial and most of their singing occurs after pair formation. Consequently, song in such species is considered to have its primary function within established pairs and in social groups (Loning et al., 2024). Also, the very clear individual distinctiveness of the single individual zebra finch song motif appears almost unique among songbirds and is likely to facilitate establishing and maintaining social bonds in their dynamic social organisation. Moreover, the difference in peak singing times at breeding colonies and social hotspots suggest behavioural routines which may allow individuals to predict when to meet whom, or how many others, at which location. This pattern might have been specific to the breeding period, as overall birds may visit hotspots more frequently when they are not tied to regular visits to a breeding site. While we cannot rule out with our data that some males may have sung at dawn at a remote roosting nest, any such possibility seems unlikely to be a wider pattern, as roosting nests typically are in the breeding colony, often near the breeding nest (Zann, 1996). Thus, dawn song there should have been picked up by the recorders. Moreover, although this remains to be studied, it seems likely that the difference in song activity between breeding sites and social hotspots might be even larger during non-breeding periods when individuals have more time available for socialising.

Next to a higher singing activity, social hotspots also included larger song networks, with more distinct individuals and singing associations than in the breeding sub-colonies, further supporting the idea that social hotspots serve as meeting places (Loning et al., 2023a). The larger hotspot network could facilitate enhanced social information exchange across familiar and unfamiliar individuals, which can be valuable in the relatively harsh and unpredictable environment zebra finches inhabit (Rafacz & Templeton, 2003; Rubenstein & Lovette, 2007). One benefit could be increased information on the breeding status of other individuals, which could inform breeding synchronisation (Mariette & Griffith, 2012b; Brandl et al., 2021) as males also sing more in groups during breeding periods than in periods without breeding (Loning et al., 2023b). While individuals can obtain breeding information for synchronisation by prospecting nests of others (Mariette & Griffith, 2012a; Brandl et al., 2019b), social hotspots could provide access to information from a wider area and in a more centralised, readily available location. Moreover, in captive zebra finches, colony sounds were found to increase breeding synchronisation (Waas et al., 2005), and the vocalisations at hotspots could have a similar effect.

While the existence of social hotspots has been shown earlier (Loning et al., 2023a), our finding that at least some individuals repeatedly were identified to associate, expands on these earlier findings. Our results here show that the hotspots are not just anonymous gathering sites but reveal a social structure with repeated encounters. The combination of encountering known and unknown individuals would allow for different levels of information exchange that can be relevant for various movement, foraging and breeding decisions in such an unpredictable environment (Benskin et al., 2002; Aplin et al., 2012; Mariette & Griffith, 2013). Zebra finch song output reflects not only, for instance, breeding stage (Loning et al., 2023b), but also early developmental background (Holveck et al., 2008; Honarmand et al., 2015), condition (Ritschard & Brumm, 2012) and testosterone levels (Cynx et al., 2005) and may therefore signal condition, foraging success, and fitness. Any such more subtly coded information might be more readily accessible from an already familiar song by known individuals.

In addition to a potential exchange of information, other factors might influence the attendance at social hotspots and breeding sites. Gathering at hotspots with many conspecifics may reduce the risk of predation through dilution or increased vigilance (Beauchamp, 2008; Lehtonen & Jaatinen, 2016). Further, breeding state may influence social network structure, as many species, including zebra finches, associate mainly in pairs during breeding periods, but form larger groups during non-breeding periods (Griesser et al., 2009; Wang & Lu, 2014; Farine & Whitehead, 2015; Kurvers et al., 2020). This suggests that the loose structure we observed may reflect the breeding season, and networks could be more cohesive outside this period.

The networks observed in this study primarily consisted of single encounters between individuals. Only 5-11% of dyads (depending on the network type) encountered each other multiple times, and only 1-3% interacted three or more times. This suggests that birds were not preferentially associating with specific others within the short time frame of our analysis. While the different network approaches yielded different quantitative values for the associations, qualitatively they all show the same pattern of network structure with only small fractions of repeated dyadic encounters. The low rate of repeated dyadic associations in our data, therefore, points to a loose, more ephemeral social structure. Yet, most likely the dyads recorded as interacting only once actually interacted more frequently, either on different days or at other locations. This is supported by the observed relationship between total singing time and the number of detected associations, and relatively short singing times for most individuals, suggesting that additional data collection most likely would reveal a substantially larger and more connected network. In addition, the use of song as a proxy for presence has limitations, as zebra finches do not sing continuously, and both males and females additionally communicate through frequent calling. Individuals were found to sing for just over half of the time that they were actually present at a hotspot (Loning et al., 2023a), meaning that some associations likely have been missed. Combining acoustic analysis with observations, including the individually specific calls, or a different methodology using an automated radio tracking system or PIT-tags (Brandl et al., 2021; Tyson et al., 2024; Tyson et al., 2026) could provide a more complete picture of the networks. Such tracking systems would also capture female associations, which are overlooked when relying on song alone in this species, although given their close spatial association (Mariette & Griffith, 2012a; McCowan et al., 2015; Tyson et al., 2024), pair members are expected to form highly similar networks (Brandl et al., 2019a).

The two relatively separate networks at the breeding colony and the social hotspots, combined with previous findings of stable pair bonds (Griffith et al., 2010; Tyson et al., 2024), suggest that their social system may share some features of a multi-level social structure as shown in aviary conditions (Zhang et al., 2025). Such a structure would have pairs as the core unit, distinct social associations at the nest and social trees as distinct higher level social units and potentially fusing of these social units later in the season. However, other defining characteristics of a multi-level society were less apparent in this free ranging population. Individual membership within grouping levels was flexible, with few repeated associations, resulting in a relatively sparse network. Furthermore, the breeding site network was not clearly nested within the social hotspot network, as we initially predicted. This does not fit the traditional definition of a multi-level society as a hierarchical social structure (Grueter et al., 2020). Further work is needed to determine whether zebra finch societies in the wild show the temporal stability and hierarchical nesting characteristics of a multi-level society or if wild zebra finch social organisation is more consistent with a generally more fluid fission-fusion social structure (Aureli et al., 2008). It is possible that a hierarchical network becomes more evident with longer-term or finer-scale data. Meanwhile, our findings give new insights into the social structure under natural free ranging conditions of a widely used model species in lab-based studies. Given this widespread use of zebra finches as a model species in the lab, a clearer understanding of zebra finch natural social dynamics may also improve the interpretation of lab findings and inform best practices in captive housing (Griffith et al., 2017).

One strength of zebra finches as a model is their individually distinct song (Immelmann, 1968; Böhner, 1983; Zann, 1990; Hauber et al., 2010; Woodgate et al., 2012), which one can use to assign individuals. Indeed, in a nomadic species such as the zebra finch (Immelmann, 1965; Zann, 1996; Mariette & Griffith, 2012a), it is a strong advantage to be able to follow unbanded individuals in space and time using acoustic monitoring. Here, individual assignment by spectrogram comparison allowed us to observe the associations on which most of the conclusions that we draw are based. Newly encountered songs were compared to a reference song database and songs that matched were assigned to the same individual. Analogously, also the songs’ deep learning representation by BirdNET differed less within individuals (as assigned by us) than between individuals. Given the very high statistical power, however, this finding is neither surprising nor indicative of a large effect size. In fact, there was a large overlap between the distributions of cosine distances of intra- and interindividual comparisons, especially when recording conditions were less similar. This suggests that on a case-by-case basis it is not possible to validate a given individual label with high certainty using this BirdNET embedding-based approach. For this validation approach we specifically left BirdNET untrained to the task of individual classification, but future studies would benefit from training BirdNET (or other machine learning models) to aid in automated individual acoustic recognition. Individual acoustic recognition by machine learning approaches has been successful in (smaller, known) captive populations (Elie & Theunissen, 2018). It remains to be seen how machine learning models perform on the task of individual recognition in the wild in this species, where recordings have much more environmental noise and the number of present individuals (classes) is unknown.

To summarise, our findings show that wild zebra finches sing near breeding sites and at social hotspots at different times of day, with a distinct lack of dawn song at the nest and the social hotspots. This highlights that the context and potential function of their singing differ from those of classically studied territorial songbirds, as previously noted for other song characteristics (Loning et al., 2024). Our establishment of using BirdNET embeddings for the validation of individual classification demonstrates a promising approach to scale up the study of communication networks, opening the door for more advanced future studies. Male song revealed different levels of social organisation, with individuals connecting with more and different conspecifics at social hotspots than at the breeding colony. A few individuals maintaining associations across both sites, can be considered as sub-units, reflecting elements of multi-level societies where information can flow through vocal interactions. However, the overall low number of associations between male dyads suggests that birds mostly move around alone or as pairs, meeting others opportunistically at social hotspots or the breeding area. This reflects a social organisation that is at the interface between a multi-level society of stable hierarchical subgroups and a more dynamic fission-fusion society with individualistic network structures. Such a dynamic system might be adaptive in the harsh and unpredictable environment of the Australian Outback, as birds have stable social subunits (the pair) connected to different familiar individuals at different locations. Encountering a variety of other, socially less stable connected individuals will allow both males and females to draw information from a wider network. This may be important to gather population-wide information, for instance, on dispersal and immigration patterns, as well as potential novel food sources, such as when unfamiliar individuals are in better condition.

Taken together, the communal singing pattern of zebra finches leading to communication networks, along with the individual distinctiveness of the song and other differences in singing compared to territorial species, widens our view on the function of birdsong. The approaches that we used here provide an example to study how information is shared across communication networks and wider social organisations in species which can be individually recognised based on their signals.

